# Spatiotemporal organization of prefrontal norepinephrine influences neuronal activity

**DOI:** 10.1101/2023.06.09.544191

**Authors:** Samira Glaeser-Khan, Neil K. Savalia, Jianna Cressy, Jiesi Feng, Yulong Li, Alex C. Kwan, Alfred P. Kaye

## Abstract

Norepinephrine (NE), a neuromodulator released by locus coeruleus neurons throughout cortex, influences arousal and learning through extra-synaptic vesicle exocytosis. While NE within cortical regions has been viewed as a homogenous field, recent studies have demonstrated heterogeneous axonal dynamics and advances in GPCR-based fluorescent sensors permit direct observation of the local dynamics of NE at cellular scale. To investigate how the spatiotemporal dynamics of NE release in the PFC affect neuronal firing, we employed in-vivo two-photon imaging of layer 2/3 of PFC in order to observe fine-scale neuronal calcium and NE dynamics concurrently. We found that local and global NE fields can decouple from one another, providing a substrate for local NE spatiotemporal activity patterns. Optic flow analysis revealed putative release and reuptake events which can occur at the same location, albeit at different times, indicating the potential to create a heterogeneous NE field. Utilizing generalized linear models, we demonstrated that cellular Ca2+ fluctuations are influenced by both the local and global NE field. However, during periods of local/global NE field decoupling, the local field drives cell firing dynamics rather than the global field. These findings underscore the significance of localized, phasic NE fluctuations for structuring cell firing, which may provide local neuromodulatory control of cortical activity.

## Introduction

Norepinephrine (NE) is a neuromodulator which plays a key role in arousal^1–3^, attention, and learning^4, 5^. Temporally brief NE release events (phasic)^6^ may facilitate orienting^7^ to sensory stimuli and serve as a “network reset” signal^8^ for neural circuits throughout the brain. Axons from the pontine locus coeruleus (LC) release NE throughout the cerebral cortex, and LC neurons can operate synchronously during sleep^3^. However, recent investigations of the anatomy of LC neurons and their dynamics have shown that they may carry distinct signals to different brain regions^4, 5^.

Structurally, LC_NE_ axons release NE sparsely from distinct varicosities^9, 10^–bulging regions of the axon where NE vesicles can be released into the extracellular space. These axonal varicosities vary in density within the cortex, with frontal cortical areas having the highest density^9^. The spatial and temporal organization of release from catecholamine axonal varicosities has been best studied in dopamine release in the striatum^11, 12^. In that context, only ∼20% of varicosities are active, suggesting spatially sparse release. Release is predominantly extrasynaptic and can be induced either by dopamine neuron action potentials or by local activation of nicotinic acetylcholine receptors on the axons of dopamine neurons^13^. Studies using false fluorescent neurotransmitters, which allow the imaging of neuromodulator release and reuptake, have revealed that NE axons function similarly, with a population of “silent” varicosities which do not release NE^14^. Thus, variation in distance from NE release sites might induce fine-scale fluctuations in NE within a brain region, although this has not been proven.

Local hotspots of NE release have been hypothesized to play a critical role in attention^15^. Indirect evidence exists that glutamate in the prefrontal cortex leads to regional NE release^16^. Moreover, pharmacological block of NE receptors in cortex hyperpolarizes neurons^17^ and diminishes their responsivity to arousal and sensory stimulation. Still, it is unknown if local fluctuations in NE release tune neuronal responses in a spatially restricted manner. NE release at this local scale could enable selective gain changes across neurons^6^, or could contribute to cell-type specific control of neural activity^18^. However, local release of NE at cellular scale has not yet been demonstrated in vivo. Thus, we sought to investigate the dynamics of local NE release within prefrontal cortex at a cellular scale.

Here, we used two-photon imaging of fluorescent biosensors for NE^19^ (GRAB-NE2h) to identify spatiotemporal scales of extracellular NE concentration dynamics in the mouse medial prefrontal cortex (mPFC). We adapted optic flow analysis^20^ to the problem of identifying patterns of NE dynamics to evaluate local NE structure. Our optic imaging preparation also included simultaneous measurement of cellular activity via a red-shifted Ca^2+^ indicator in mPFC neurons (jRGECO1a). Generalized linear modeling defined the relationship between NE and mPFC Ca^2+^ activity. Finally, pharmacological tools were used to causally disrupt this relationship between NE and mPFC neural activity. These optic approaches to define neuromodulator dynamics in vivo may facilitate further studies of the fine spatial scales of neuromodulation.

## Methods

### Animals

Adult male mice (age 8-12 weeks at the start of experimental procedures) of the C57BL/6J strain (Stock No. 000664, Jackson Laboratory) were used. Mice were group housed (3–5 mice per cage) on a 12:12-h light/dark cycle with free access to food and water. All imaging experiments were performed during the light cycle. All experimental procedures were approved by the Institutional Animal Care and Use Committee at Yale University.

### Light sheet microscopy

Transgenic mice positive for DBH-TdTomato (Ai9, The Jackson Laboratory) were perfused using 10 mL 1X PBS + 10 U/mL heparin (Millipore Sigma, H3393-50KU) followed by 10 mL 4% paraformaldehyde. The brains were then isolated, stored in 13 mL of 4% paraformaldehyde at 4°C overnight with gentle shaking, and then rinsed with 1X PBS. The brains were then shipped to LifeCanvas Technologies (MA) in 1X PBS +0.02% sodium azide for whole brain imaging. Whole mouse brains were processed using the SHIELD protocol (LifeCanvas Technologies). Samples were cleared for 1 day at 42 °C and then actively immunolabeled. Each sample was labeled with mouse anti-mouse NET (IGg2b, clone 2-3B2) followed by fluorescently conjugated secondary antibodies. Samples were imaged at 3.6X using a SmartSPIM axially-swept light sheet microscope (LifeCanvas Technologies). Images were tile corrected, de-stripped, and registered to the Allen Brain Atlas (Allen Institute).^21^

### Two-Photon imaging

#### Surgery

All surgical procedures were performed on mice placed in a stereotaxic apparatus (David Kopf Instruments). Adult male mice (N=3) were first anesthetized with isoflurane in oxygen (3-4% for induction, 1-1.5% for rest of surgery). The mouse laid on top of a water-circulating heating pad (Gaymar Stryker) set at a constant temperature of 38 °C. Eyes were lubricated with ophthalmic ointment. Carprofen (5 mg/kg, S.C.; 024751, Henry Schein Animal Health) and dexamethasone (3 mg/kg, i.m.; 002459, Henry Schein Animal Health) were used preoperatively to reduce pain and inflammation. The scalp was sterilized with swabs of ethanol and betadine before an incision was made to expose the skull. A ∼0.5 mm craniotomy was made using a dental drill, first for viral injections then for lens implantation ∼3 weeks after (see below). Immediately after surgery and on each of the next 3 days, carprofen (5 mg/kg, S.C.) was given to the mice. All mice recovered for at least two weeks after GRIN lens implantation before acclimating to the imaging environment.

For viral injections, we made a small skin incision and craniotomy (∼0.5 mm diameter) in the right hemisphere centered 1.7 mm anterior to bregma, and 0.3 mm lateral to midline to target the medial prefrontal cortex (prelimbic region). A glass pipette (3-000-203-G/X, Drummond Scientific) was pulled to a fine tip (P-97 Flaming/Brown Micropipette Puller, Sutter Instrument), front-filled with two adeno-associated viruses in a 1:1 mixture and then slowly lowered into the brain using the stereotaxic apparatus at 4 sites corresponding to corners of a 0.2-mm wide square centered on the above target coordinate. At a depth of 1.2 to 1.4 mm below the dura, we injected the virus using a microinjector (Nanoject II, Drummond Scientific). We injected 4.6 nL per injection pulse, with ∼10 s between each injection pulse, for a total injection volume of ∼478.4 nL split equally across injection sites. To reduce backflow of the virus, we waited at least 5 min after finishing injection at one site before retracting the pipette. The brain was kept moist with artificial cerebrospinal fluid (ACSF, in mM: 5 KCl, 5 HEPES, 135 NaCl, 1 MgCl2, 1.8 CaCl2; pH 7.3). After completing the injections, the craniotomy was covered with silicone elastomer (0318, Smooth-On, Inc.), and the scalp sutured (Henry Schein).

Three weeks following the viral injection, mice were given a second surgery to implant a gradient refractive index (GRIN) lens^22^ (Inscopix; 1.5 mm length cylindrical lens with 0.5 mm diameter). After initial steps consistent with the viral injection surgery, an incision was made to remove all the scalp along the dorsal surface of the skull, which was then cleaned to remove connective tissues. We remade a 0.5-mm diameter craniotomy over the previously targeted location. The GRIN lens has a working distance of ∼0.1 mm, so the bottom of GRIN lens was targeted to 0.1 mm dorsal to the center of the prior virus injections (i.e., 1.2 mm below the dura). To prepare the cortex overlying the target region for mass effect due to the GRIN lens, a sterile 30-gauge syringe was slowly lowered into the implant’s target position at a rate of 0.1 mm per minute. The syringe was held in the implant’s target position for 30 minutes then slowly retracted. The GRIN lens was then lowered into the target implant location at the same rate as the syringe prior. A high-viscosity adhesive (Loctite 454) was applied around the bottom rim of the GRIN lens collar to create a seal between the lens shaft and the skull. After the adhesive cured, a custom-made stainless steel headplate (eMachineShop.com) was secured to the skull using quick adhesive cement (C&B Metabond, Parkell).

#### Viruses

To image Ca^2+^ transients in pyramidal neurons along with local fluctuations in norepinephrine concentrations, we injected AAV1-Syn-NES-jRGECO1a.WPRE.SV40 (GENIE project)^23^ at a titer of 1×10^13^ genome copies (GC) per milliliter alongside AAV9-hSyn-GRAB_NE2h_ at a titer of ∼1×10^13^ (Li lab)^19^. Prior to virus injection, a 1 μL solution containing a 1:1 mixture of the two constructs was loaded into the microinjection pipette. A total of 478.4 nL of virus was injected to prelimbic cortex. The GRAB-NE2h sensor is a mutated alpha-2a adrenergic receptor tagged with an intracellular GFP that fluoresces upon NE binding to the extracellular domain and has similar affinity for NE as the endogenous alpha-2a receptor.

#### Two photon acquisition

Two-photon imaging was performed with a laser-scanning microscope (Movable Objective Microscope, Sutter Instrument) using a tunable TI:Sapphire femtosecond laser excitation source (Chameleon Ultra II, Coherent) controlled by ScanImage software^24^. Laser scanning was performed using a 980 nm excitation beam focused through a water immersion objective (XLUMPLFLN, x20/0.95 NA, Olympus) incident on an implanted GRIN lens with time-average laser power less than 120 mW during data acquisition. Fluorescence emissions were separated into red (jRGECO1a) and green (GRAB-NE2h) channels using filters (Semrock and Chroma) with center emission wavelengths 605 nm and 525 nm, respectively. Emitted photons were collected by GaAsP photomultiplier tubes. Planar images were acquired at 256 x 256 pixels at 1.47 or 0.74 μm per pixel resolution using bidirectional scanning with a 30.02-30.08 Hz frame rate. Mice habituated to head fixation under the two-photon microscope for 3-5 days of increasing duration prior to experimental data collection. Head fixed mice stood on a spinnable disk platform, which allowed volitional forward and reverse locomotion under the microscope while limiting three-dimensional head displacement. Mice underwent a baseline imaging session before the experiment to locate a field of view within the GRIN lens focal plane with the most jRGECO1a-positive somata flanked by GRAB-NE2h-positive signal, which primarily expressed on neuropil. The plane identified during the baseline session was used for all subsequent imaging sessions on a per mouse basis. For experimental imaging, mice were collected from their home cage and given an injection (i.p.) of saline (10 ml/kg) or desipramine (10 mg/kg), then situated under the two-photon microscope for a 15-20 minute imaging session. The latency between injection and the imaging session was timed out to 10 minutes for consistency between mice.

#### Drug treatment

On subsequent days, mice were imaged 10 minutes after injection with either saline or a NE transporter inhibitor (desipramine, 10 mg/kg I.P.), which induced gradually increasing NE levels across the imaging region. The injection conditions were counterbalanced across mice. Since NE accumulates everywhere simultaneously in the absence of reuptake, the importance of phasic NE release is diminished following the desipramine treatment condition.

#### Initial data processing

Two photon imaging data for each channel were separated then median filtered within pixel using a three-frame window, and downsampled in time using mean binning to a final frame of ∼15 Hz. Each channel’s fluorescence data was motion corrected for translations by non-rigid body registration using the NoRMCorre toolbox implemented in MATLAB^25^. Each frame of imaging was realigned to the temporal average collected over the first five minutes of the experiment. The merged two-color two-photon images revealed an expression pattern whereby putative somata tended to show high jRGECO1a signal flanked by background neuropil showing high GRAB-NE2h signal. Thus, we approached subsequent analysis by first selecting putative cell bodies (i.e., locations with high jRGECO1a expression) as regions of interest (ROIs) using a custom graphical user interface in MATLAB. Putative somatic ROI masks were manually traced for each mouse using the temporal average from the red fluorescence channel. A total of 83 putative cell bodies were selected across all mice. For each putative cell body, the fluorescence timecourse was extracted as the pixel-wise average fluctuations within the ROI mask. We then operationalized the meshwork of neuropil-based GRAB-NE2h signal as an NE “field.” Each putative cell body ROI was used to extract a cell-adjacent “local” NE field: a ∼15-20 μm diameter annulus centered on the ROI but excluding the ROI itself. Further, each ROI was assigned a corresponding “global” NE field, defined as all pixels in the field of view excluding the cell body and its local NE field.

#### Accounting for bleed-through in two-photon imaging

Bleed-through (also called channel cross-talk) in two-photon imaging is when some fluorescence from one fluorophore is registered by the microscope as a signal from the other fluorophore.^26^ Thus, there may be mixing of the two signals. We corrected for bleed-through by taking the residuals of the correlation between both channels at each pixel. The channel of interest (either jRGECO1a or GRAB-NE2h) was placed on the y-axis. The time series of the residuals, which represents the pure activity of the channel of interest, was used in further analysis in place of the raw fluorescence time series of the channel of interest.

#### Calculating the spatial correlation of the GRABNE signal

The spatial correlation of the GRAB-NE2h signal was calculated by dividing the field of view into a grid of 10 μm by 10 μm squares. For each given square, the Pearson’s correlation comparing the GRAB-NE2h fluctuation within the square to that of every other square was calculated. The resulting vector the correlation values were ordered by distance to the original given square.

### Optic Flow

#### Generating quiver plots and extracting patterns

To measure NE release and reuptake, we performed an optic flow analysis (Horn-Schunk method) on pseudo-colored fluorescence recordings of the GRAB-NE2h sensor, with high fluorescence regions (fluorescence above the mean) depicted yellow and low fluorescence regions (fluorescence below the mean) depicted blue. Before performing optic flow, the fluorescence recording was first spatially gauss-filtered using a sigma value of 12. The data was then temporally smoothed using a 5-second kernel, detrended, and z-scored with respect to time. Optic flow was then performed using the Horn-Schunk method, which allows us to estimate small motion between consecutive frames of our fluorescence recordings, making it a good analysis to track small-scale phasic fluctuations in NE signal. The results of the optic flow analysis were a series of motion vector plots for each frame of our recordings. The time and central location of different patterns were extracted from these motion vector plots using the findAllPatterns function in the NeuroPatt Toolbox (https://github.com/BrainDynamicsUSYD/NeuroPattToolbox) and the calc_save_SourceSink function from the OFAMM toolbox (https://github.com/navvab-afrashteh/OFAMM). Both toolboxes extracted sources, sinks, saddles, spiral in, and spiral out patterns (if present).

#### Extracting NE release and reuptake events

We observed that regions of both high and low fluorescence appear and disappear in a pattern of first expansion (a source) and then recession (a sink). Sources and sinks extracted by pattern detection (see NeuroPatt Toolbox above) were further categorized by the fluorescence value at the center of the pattern (either greater than one standard deviation above mean fluorescence or less than 1 standard deviation below mean fluorescence). Growing regions (sources) were taken as the onset of a NE event (either release or reuptake), while shrinking regions (sinks) were taken as the offset of a NE event. Together, a high fluorescence source (onset of release) followed by a high fluorescence sink (offset of release) represent one release event, the time and location of which is given by pattern detection algorithms. Similarly, a low fluorescence source (onset of reuptake) followed by a low fluorescence sink (offset of reuptake) represent one reuptake event.

#### Calculating the spatial and temporal correlations for release and reuptake sites

Spatial correlations for release and reuptake were calculated by linearizing heatmaps of the locations for all release or reuptake events that occurred in a run. These matrices were then linearized and used to calculate the spatial correlations between different patterns. The temporal correlation between patterns was calculated using the time series of the total number of release or reuptake sites at each time point.

#### Predicting NE synchrony using time and location of release sites

NE synchrony is defined as the sliding correlation between local NE and global NE. The sliding window used is 30 seconds. This time series shows the periods when local and global NE become decorrelated. Each local field has its own unique timeseries of NE synchrony. For each local field, we also calculated a “nearest distance to release” time series, defined as the nearest distance to release for a given local field at each time point. We then fitted a logistic regression predicting NE synchrony using distance to release.

#### Accounting for heterogeneity in expression of the sensor

There is some concern that the spatial correlations of patterns extracted by optic flow may be artificially inflated due to the underlying spatial expression pattern of GRAB-NE2h expression. Specifically, there is a slight correlation between the spatial location of any pattern and the raw fluorescence level of GRAB-NE2h, suggesting locations with greater NE fluctuations occur where there is greater sensor expression. To account for this, we took the correlation between activity patterns with their underlying correlation to the fluorescence level of the GRAB-NE2h signal regressed out.

#### Controlling for the potential effects of processed noise

Before performing the optic flow analysis, our dataset was spatially and temporally smoothed with a 4-micron spatial gauss filter and a 5-second temporal filter. We generated 500 random datasets and calculated the spatial and temporal correlations of “release” and “reuptake” sites (see “*Extracting NE release and reuptake events*”) amongst other patterns (e.g., “spiral-in,” “spiral-out”). The random datasets were computer-generated random time series that had the same size, mean across time, distribution, and standard deviation across time as our experimental datasets. Using the correlation distributions from the 500 randomized datasets, the observed correlations from the true dataset were examined for significance using a two-tailed test (*P* < 0.05).

### Models predicting cell Ca^2+^ activity

#### Predictors used in the GLM

We fit generalized linear models predicting neuronal Ca^2+^ dynamics^27^. Independent predictors were local NE, global NE, global Ca^2+^ dynamics, and the interaction between local NE and global NE (see below). The time points were then further separated into high NE synchrony intervals and low NE synchrony intervals.

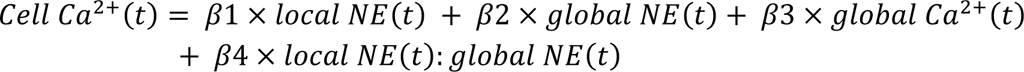

#### Cell Ca2+ dynamics (response variable)

the jRGECO1a sensor fluctuations within one putative cell body, averaged over space to yield a single fluorescence timeseries. The timeseries is then detrended, temporally smoothed with a 1-second kernel, and z-scored with respect to time.

Global Ca^2+^ dynamics (predictor): The mean jRGECO1a sensor fluctuations in all putative cell bodies within the field of view minus the cell body of interest (i.e., response variable). The timeseries is again averaged across space, detrended, temporally smoothed with a 1-second kernel, and z-scored with respect to time.

#### Local NE (predictor)

The GRAB-NE2h sensor fluctuations within a radius of 15 microns of the cell body excluding the area of the cell body itself. The GRAB-NE2h fluctuations are averaged over space to yield a single fluorescence time series. The time series is then detrended, temporally smoothed with a 1-second kernel, and z-scored with respect to time.

#### Global NE (predictor)

The mean GRAB-NE2h sensor fluctuations within the entire field of view excluding the local NE field and the cell body of interest. The timeseries is averaged over space, detrended, spatially smoothed with a 1-second kernel, and z-scored with respect to time.

#### Average NE synchrony (used to split time points)

NE synchrony is defined as the sliding correlation (computed with a 30-second window) between a given local NE field and its global field (as defined above). The NE synchrony time series for all cells are highly coupled, indicating global periods of high NE synchrony and global periods of low NE synchrony. The average NE synchrony time series for each run is used to divide all time points into intervals of high NE synchrony and low NE synchrony.

It is important to note that each putative cell body has its own unique local NE field, global NE field, and global Ca^2+^ timeseries. Thus, separate GLMs were fitted to predict the Ca^2+^ dynamics of each putative cell body. The GLMs were fit using forced entry, so each model included all predictors.

#### Models predicting differences in mean correlations between conditions

Pairwise correlations between local NE fields (local/local NE correlations) and correlations between the local NE field and corresponding global field (local/global NE correlations) were compared between conditions using a linear mixed effects model (LME model). For each type of correlation, the drug treatment (either saline or desipramine) was included in the model as a main effect, while the effect of cell number and mouse were included as uncorrelated random effects (see equation below). The LME model was used to account for the fact that multiple observations were included from the same cell and mouse. The LME model was fit using the Matlab ‘fitlme’ function, which uses a trust region based quasi-Newton optimizer.

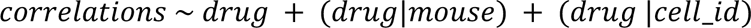

## Results

To understand how spatiotemporal dynamics of norepinephrine (NE) release in the prefrontal cortex affect neuronal firing, a green NE sensor (GRAB-NE2h) and a red calcium sensor (jRGECO1a) were expressed pan-neuronally using AAV transfection (AAV1-hSyn). Two-photon imaging of fine-scale spatiotemporal neuronal and NE dynamics was subsequently carried out using an inserted gradient-refractive index (GRIN) lens across layer 2/3 of prelimbic medial prefrontal cortex during spontaneous locomotion. A total of 83 putative neurons (N = 3 mice) were thus recorded while measuring local NE dynamics using this approach. Mice were allowed to run freely on a spinning disc treadmill while cellular resolution NE and Ca^2+^ dynamics were recorded optically. On subsequent days, mice were repeatedly imaged 10 minutes after injection with either saline or a NE transporter inhibitor (desipramine, 10 mg/kg i.p.), which induced gradually increasing NE levels across the imaging region. Modeling of the relationship between neuronal firing and local norepinephrine was performed using generalized linear models (GLM) to determine the relationship between NE and Ca^2+^ in neurons across spatial scales. Optic flow analysis was used to delineate patterns of NE release and reuptake in mPFC and to relate these patterns to the ongoing activity of neurons.

### Light-sheet microscopy reveals high NE axon density and low transport density in mPFC

We hypothesized that spatiotemporal patterns of NE release would be strongly influenced by both the density of NE axon innervation as well as the density of the NE transporter^28^, which removes extracellular NE. Specifically, a low ratio of neurotransmitter release to reuptake leads to highly localized neurotransmitter release events^29^ (such as synaptic glutamate), whereas a high ratio might lead to events over diffuse spatiotemporal scales (e.g., extrasynaptic volume transmission^28^. These larger spatial release events might be well suited to measurement using extracellular fluorescent biosensors. Thus, we conducted whole brain light-sheet microscopy^21^ of transgenic mice expressing fluorophore in NE axons (DBH-Cre x flex-TdTomato mice) after tissue clearing and NET antibody staining to identify regions with a high density of NE axons from the locus coeruleus (LC) but low density of NET (Figure 1A). We found that the density of LC innervation in the mPFC is elevated compared to the visual and somatosensory cortex (2-sample t-test, *P* = 2.4913e-08) (Figure 1B). The density of NET has been previously shown to be lower in cortex than in some subcortical structures^30^. These combined findings suggest NE release and reuptake in mPFC may be subject to diffuse, extrasynaptic spatiotemporal dynamics which could be resolved via *in vivo* two-photon imaging.

**Figure 1.**
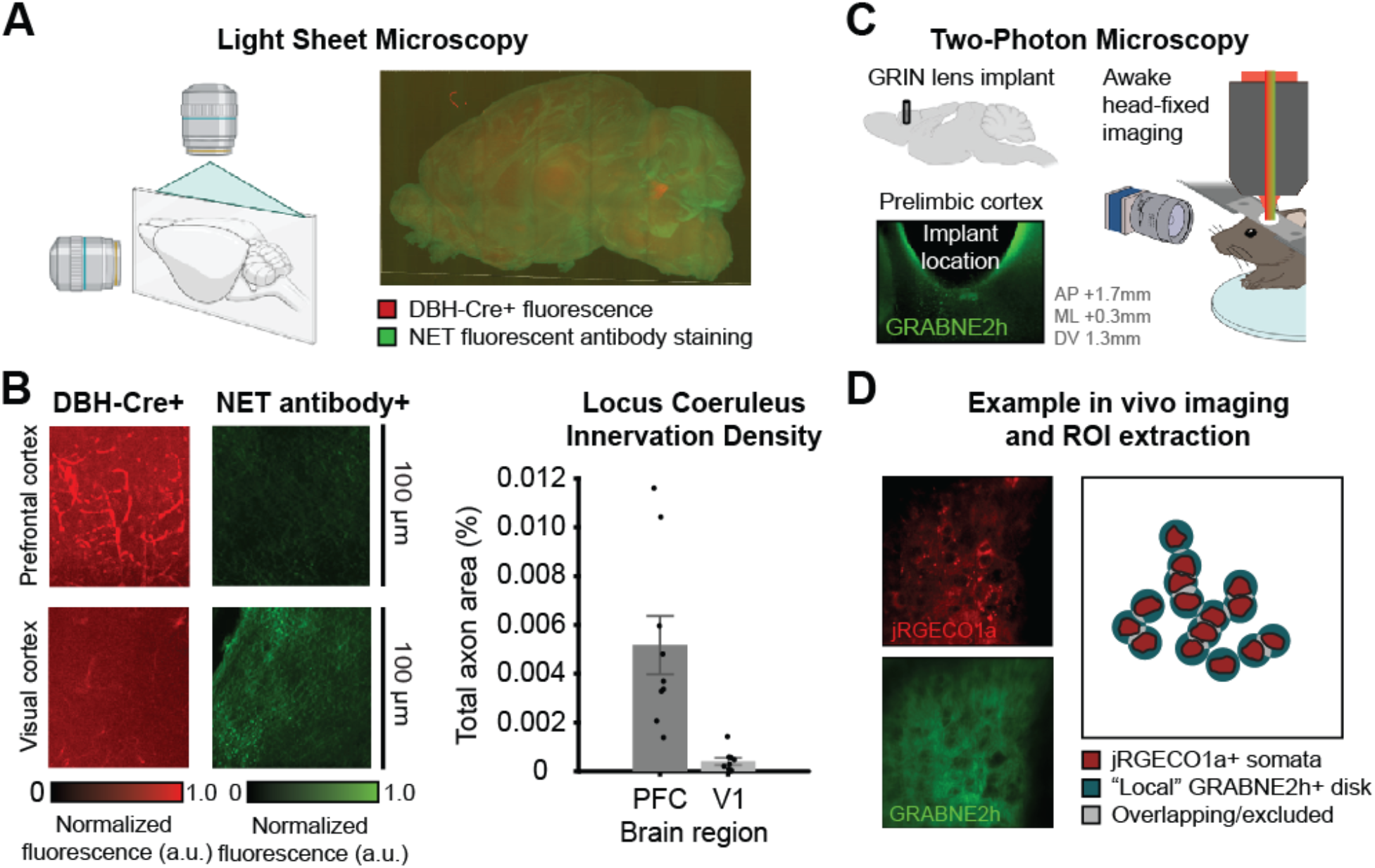
Unique anatomy in the PFC allows for observation of spatially diffuse NE. **A)** Light sheet microscopy was performed by fixing the brain of DBH-TdTomato mice, staining with an anti-NET antibody, and subsequently imaging horizontal slices. Light sheet microscopy 3D images of NET (green) and LC axons (red). **B)** PFC receives denser LC innervation than the visual cortex (left) but has a lower density of NET (center). Comparison of axon density in the PFC vs. the visual cortex from n = 10 brain slices (right). **C)** Mice were infected with two adeno-associated viruses to express jRGECO1a (a red Ca^2+^ sensor) and GRAB-NE2h (a green NE sensor). Simultaneous Ca^2+^ and NE dynamics were measured in awake head-fixed mice using two-photon imaging through a GRIN lens implant targeting medial PFC. **D)** An example field of view from two-photon imaging of jREGCO1a (top) and GRABNE2h (bottom). Ca^2+^ dynamics were extracted as the average fluctuations in manually-traced putative cell body regions of interest. Each putative cell body was assigned its own local NE field, defined as the 15 μm circular disk surrounding the somatic region of interest (excluding pixels assigned to the cell body itself and any overlapping NE fields).

### Two-photon imaging of spatiotemporal dynamics

To measure NE dynamics and their influence on cellular firing in vivo, we used dual-color two-photon imaging of mPFC to simultaneously measure fluctuations of red Ca^2+^ sensor (jRGECO1a^23^) and a green NE sensor (GRAB-NE2h^19^). jRGECO1a provides a readout of intracellular Ca^2+^ as a proxy for neuronal activity, whereas GRAB-NE2h is a genetically modified NE receptor that senses the extracellular concentration of NE. We expressed both jRGECO1a and GRAB-NE2h under the control of the pan-neuronal hSyn promotor using a dual virus approach and were able to obtain expression of both sensors. Since the mPFC is located beyond the scattering limit of standard two-photon imaging through a cranial window, these sensors were visualized (Figure 1D) using an implanted gradient refractive index (GRIN) lens. Since light traveling through GRIN lenses may suffer from chromatic aberration, we used a single wavelength (980 nm) to excite both green and red fluorophores thus ensuring that the imaged red and green signals originated from the same spatial location. GRIN lens two-photon imaging of neural ensembles and NE fields with a single wavelength of excitation and spectrally distinct emission wavelengths thus permitted us to identify spatiotemporal dynamics of these two sensors in a deep brain structure (mPFC).

### NE fluctuations have local structure that transiently decouples from global NE fluctuations

Neuromodulators can elicit their effects on neural excitability and plasticity either through synaptic release or volume transmission. NE release has recently been shown to vary across brain regions^5^ and we hypothesized that a two-photon imaging approach would facilitate the observation of variation in NE spatial dynamics at a cellular scale. Within the mPFC, NE is released at discrete sparse axonal varicosities of LC_NE_ neurons. Spatiotemporal dynamics of NE could be influenced by diffusive spread of NE^28^ from these varicosities of distinct LC_NE_ neurons firing asynchronously, or by local regulation of varicosities from the same LC_NE_ neuron. GRAB-NE2h is expressed diffusely, providing a measure of extracellular NE dynamics at each position in the field of view. By using a circular area around each putative neuron, we sought to obtain a local NE signal which could be compared with the average global NE across the entire remaining field of view (Fig. 2A). We hypothesized that local NE signals might be asynchronous and vary with distance from release site. Therefore, we first measured the correlations between local NE regions as well as the correlations between local NE regions and the global NE signal (Fig. 2B). We observed that temporal correlations between local NE sites were moderate, but were significantly less than those between local NE and global NE (linear mixed effects models showed a decrease of 0.13117, *P* = 2.5904e-07) indicating temporal heterogeneity in local NE fields. We next sought to determine if, in keeping with the diffusive spread hypothesis, local NE correlations showed a characteristic spatial extent. Spatial autocorrelation of NE was determined using 10 μm patches in a grid (see Methods) and was approximated well by a double exponential model, indicating that local NE correlations fall off with spatial distance (Fig. 2C). These dynamics thus appeared compatible with a model in which diffusive spread of NE from release sites contributes to variation in local NE dynamics.

**Figure 2.**
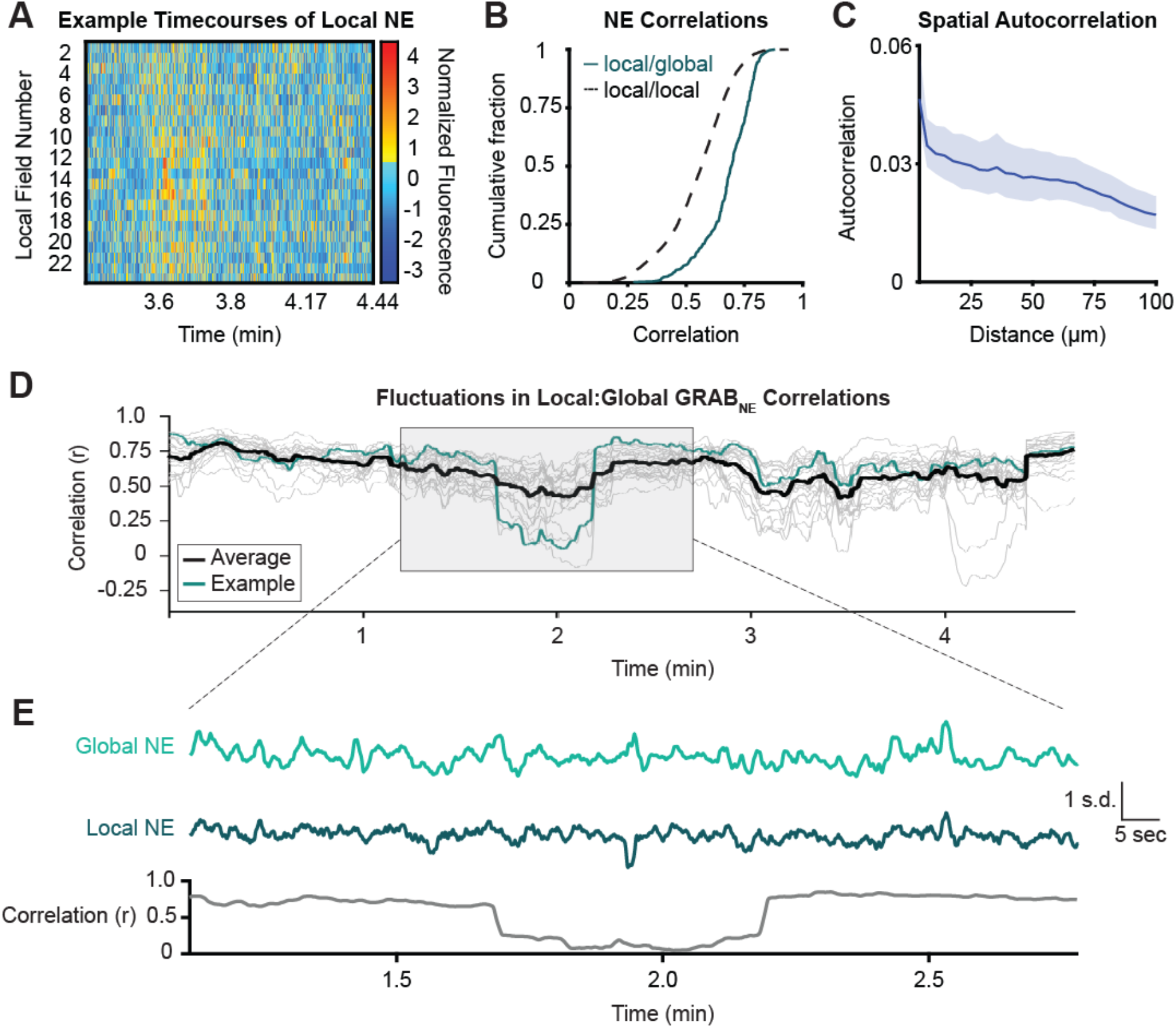
Local norepinephrine has a characteristic spatial and temporal scale. **A)** Example of movement-corrected local (15 mm) NE fluorescence around putative cells. Different local NE fields peak at different times. **B)** Cumulative distribution function (CDF) showing significantly lowered correlations between local NE fields to each other (dotted) compared to the correlations between local NE fields and the global NE fluorescence (solid) (KS test p = 2.9983e-57). **C)** Spatial autocorrelation function of NE fluorescence: temporal correlation of patches (10 μm^2^) of local NE fluorescence to other patches as a function of distance **D-E)** Example traces of NE synchrony time series (local NE:global NE sliding correlation with a 30 s window) over (D) 5 minutes for 28 local NE regions (average in black, example synchrony trace from 1 local NE field in blue) (E) 1 minute corresponding to shaded area of (D). Transient dips in correlation are observed when the local and global NE fluorescence become decoupled.

Prior work has demonstrated that LC neurons and axons fire synchronously to aversive events^4, 5^, but that axons can be asynchronous for appetitive events^5^. We hypothesized that these distinct patterns of LC_NE_ neuronal activity (local asynchronous and global synchronous) might be reflected at cellular scale within mPFC in the temporal correlations of local and global NE fields. Release at local NE sites could vary over time, either due to local regulation of release^13, 31^ or due to variation in the activity of specific NE neurons^4, 32^. We found that the local NE field and the global NE field are overall positively correlated and are more correlated than the local NE fields are to each other (Figure 2B). The correlation of NE within pairs of local regions diminished with distance (Fig. 2C). However, the correlation structure between the local and global field was not stable over time – there were alternating periods of high and low NE synchrony, as measured by the sliding local:global NE correlation (Fig. 2D-E).

### NE locally influences neuronal activity

Since the local NE dynamics around neurons differ from one another, we hypothesized that this may influence neuronal activity on a local spatial scale. To elucidate the relationship between local NE and cell firing, we fit generalized linear models predicting each individual cell’s Ca^2+^ fluorescence using its local NE, the average remaining global NE, their interaction, and the average remaining bulk Ca^2+^ fluorescence (Figure 3A). This modeling approach enabled us to determine whether local NE was predictive of cellular activity, and whether that relationship could be explained by other variables such as shared variability between cells (i.e., neuropil Ca^2+^ fluorescence). We fit a separate model for each cell and then examined the resulting distributions of beta weights for each predictor to identify consistent predictors of neuronal activity. Models were also separately fit during times of low and high NE synchrony, as we hypothesized that local NE might have a greater influence on Ca^2+^ fluorescence during periods of decoupling from the global NE signal (low NE synchrony). Using all time points or only high NE synchrony time points, all predictors are significantly positive (Fig. 3C). The significance of the interaction term confirms that local NE and global NE are highly coupled for high synchrony time points. However, when only low synchrony time points are used, only local NE (mean beta = 0.0773, 1-sample *t*-test, *P* = 8.4224e-31) and global Ca^2+^ (mean beta = 0.2941, 1 sample *t*-test, *P* = 1.1321e-47) are significant predictors of cell Ca^2+^ (Fig. 3D). The interaction term is not significantly positive, which confirms that local NE and global NE are decoupled for these time points. These results indicate that local NE is a better predictor of cell Ca^2+^ than global NE and decouples from global NE during low synchrony periods.

**Figure 3.**
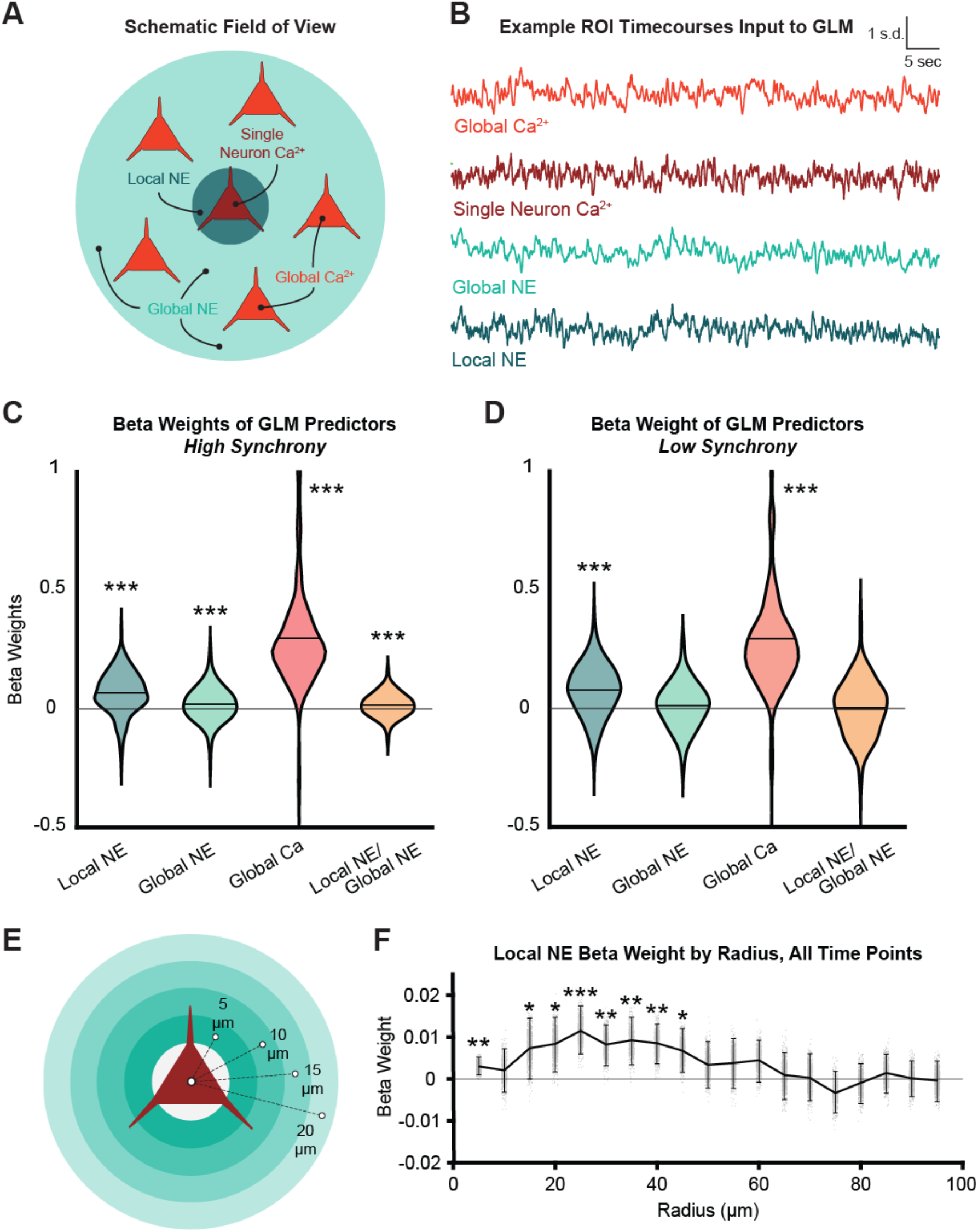
Local NE predicts cellular Ca^2+^ dynamics. **A)** Schematic of variables used in generalized linear modeling. Single neuron Ca^2+^ – jRGECO1a fluorescence of one neuron (response variable); Global Ca^2+^ - jRGECO1a fluorescence of all other cells in the field of view (FOV); Local NE – the GRAB-NE2h fluorescence in a radius of 15 μm around the target neuron (but not including the neuron pixels); Global NE – the GRAB-NE2h fluorescence in the FOV (but not including the chosen local NE pixels). **B)** Example time series of all inputs and response variables for GLM modeling. NE synchrony is the sliding correlation with a 30s window between local NE and global NE. NE synchrony is used to select time points for modeling (separate models are fitted for high and low synchrony periods). **C)** GLM models were individually fitted to N = 326 ROI Ca^2+^ fluorescence separately for periods of high and low synchrony (as in Figure 2E). During high synchrony, the results demonstrate significant separate correlation of local NE, global NE, global Ca^2+^ as well as local:global NE interaction. **D)** Global NE and local:global NE interaction terms were reduced to a non-significance during low synchrony relative to high synchrony periods. **E**) Schematic of repeated model fitting at expanding radii for the local NE field (5 microns – 100 microns) to determine spatial extent of local NE-cell Ca^2+^ relationship. **F)** Beta coefficient distribution (n=326 cells) as a function of expanding radii demonstrating highest peak correlation at 25 um for the low synchrony condition.

We next considered the possibility that the predictive power of local NE might relate to artifactual contamination of the green (GRAB-NE2h) channel from the red (jRGECO1a) channel within the same spatial region. Calcium imaging fluorescence within cellular ROIs is known to be correlated with the dynamics in surrounding local areas due to the activity of dendritic processes^33^, representing a potential confounding variable. All GLM analyses were conducted by first regressing out the GRAB-NE2h or the jRGECO1a signal from the pixels within each local NE or neuronal region, respectively, to ensure that potential light leak across fluorescence channels could not be responsible for the observed local NE-cell relationship. Comparing the local NE-cellular Ca^2+^ relationship before and after this regression procedure, there was no significant difference in GLM model fits (Fig. 3E), suggesting that this potential confound could not explain the observed results.

Furthermore, we reasoned that the local NE-cell relationship ought to fall off spatially with distance if it were due to detection of true spatiotemporal dynamics of NE influencing cell firing. We found that the significance of the local NE predictor peaks at 25 μm from the center of the cell body and falls off by 45 μm (Fig. 3G). Thus, we observed that local NE provided the highest predictive value for cell firing within a local area of 25 μm even after removing any correlation to the neuropil calcium signal.

### Pharmacological disruption of NE fluctuations weakens local NE-cell firing relationship

Since these analyses demonstrated a local relationship between local NE release and adjacent cell firing, we sought to determine if this relationship would be diminished after altering NE dynamics. The NE transporter removes the extracellular NE observed in our study, thus limiting the spatial scale of diffusion^28^. We administered a pharmacological inhibitor of the NET (desipramine) in order to cause synchronous accumulation of NE across the field of view^19^. Under the influence of desipramine, spatially structured NE dynamics are disrupted, as evidenced by the absence of large, slow fluctuations in local NE time series when desipramine is administered (Figure 4B). After detrending the long-term NE accumulation, both local NE/local NE as well as local NE/global NE correlations at the seconds scale are decreased with desipramine (linear mixed effects models shows a decrease of 0.10824 and 0.095292, *P* = 0.038692 and 0.0096055 respectively), indicating a loss in spatial and temporal structure. When the GLM modeling procedure is repeated (see above), we find that all predictors, including the interaction term, are significant. As in the saline condition, local NE is a significantly more positive predictor of cell firing compared to global NE. However, the mean beta weight of local NE is significantly decreased in the desipramine condition compared to saline (Figure 4E 2-sample *t*-test, *P* = 0.0125), showing that desipramine weakened the relationship between local NE and cell firing. Thus, the desipramine causal manipulation is consistent with our hypothesis and demonstrates that the relationship between local NE and cellular dynamics depends on the NE transporter.

**Figure 4.**
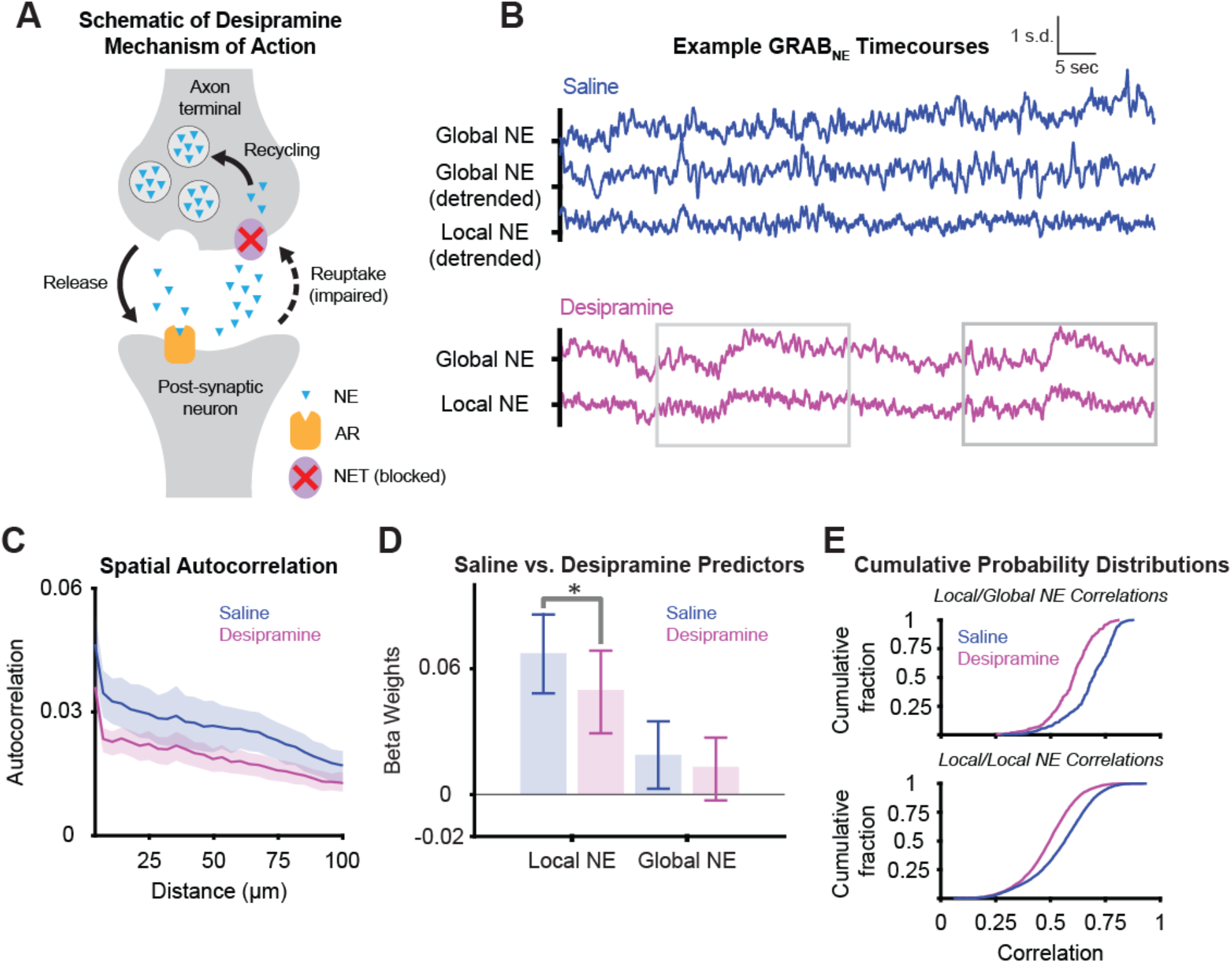
NE reuptake inhibition disrupts local NE structure. **A)** Desipramine works by blocking the NE transporter, leading to the global accumulation of extra-synaptic NE. **B)**--Representative examples of local NE during either NE transporter inhibition by desipramine (top, blue) or saline (bottom, magenta). Desipramine condition is shown both prior to and after detrending. The detrended time series are used to define spatiotemporal correlations in subsequent analyses. **C)** Desipramine and saline spatial autocorrelation function of NE fluorescence calculated as the temporal correlation of patches (10 pixels) of local NE fluorescence to other patches as a function of distance.(desipramine space constant: 0.8026 mm 95% CI: (0.2577, NR); saline space constant: 1.3430 mm 95% CI: (1.3430, 2.9283)) **D)** GLM models were individually fitted to N = 310 ROI Ca^2+^ fluorescence traces for all time points, demonstrating significantly decreased beta weights for local NE (two-sample t-test p = 0.0125), but not global NE **E)** Cumulative probability distributions (CDF) of the correlation of local NE fields to each other (top) and local NE fields to the global NE field (bottom) for desipramine and saline shows significantly correlations for desipramine (Kolmogorov–Smirnov test p = 1.9717e-101 (top) and p = 2.3206e-24 (bottom)).

### Optic flow analysis reveals release-site dependent mechanism for NE desynchrony

Optic flow analysis has been used to characterize complex spatiotemporal patterns in neural activity, including spreading waves in local field potentials^34^ and in calcium imaging^35^. We reasoned that fine-scale neurotransmitter spread via two-photon imaging of GPCR-based sensors might enable direct observation of neurochemical spatiotemporal dynamics, and their correspondence to ongoing neural activity. NE spatiotemporal dynamics may be influenced by the structure of release from spatially defined varicosities on locus coeruleus axons^10^, as well as by reuptake by NE transporter on NE axons and astrocytes^36^. Therefore, we calculated optic flow fields in the NE fluorescence channel (Figure 5A) which enabled us to determine the gradient of fluorescence in local regions from consecutive frames of two-photon imaging acquisitions (see Methods). This gradient defines the direction of NE spread at each pixel over time. These patterns of the NE field can be used to define time and location of NE spread and shrinkage (expanding or contracting NE fluorescence, Figure 5B). We observed that the same spatial location can go from being a putative reuptake site (low fluorescence; Figure 5A top right panels) to being a putative release site (high fluorescence; Figure 5A bottom right panels). NE release and reuptake tend to occur at the same location (mean spatial *r* = 0.68442, 1-sample t-test, *P* = 6.07086e-16, Fig.5C), but release and reuptake events are temporally anti-correlated (mean temporal *r* = −0.258827, 1-sample *t*-test, *P* = 0.0005632, Fig 5C). The distance to the nearest release site for each time point (Figure 5E-F) predicted the local-global NE synchrony, with more distance from release sites leading to a more homogeneous NE field (greater local-global synchrony).

**Figure 5.**
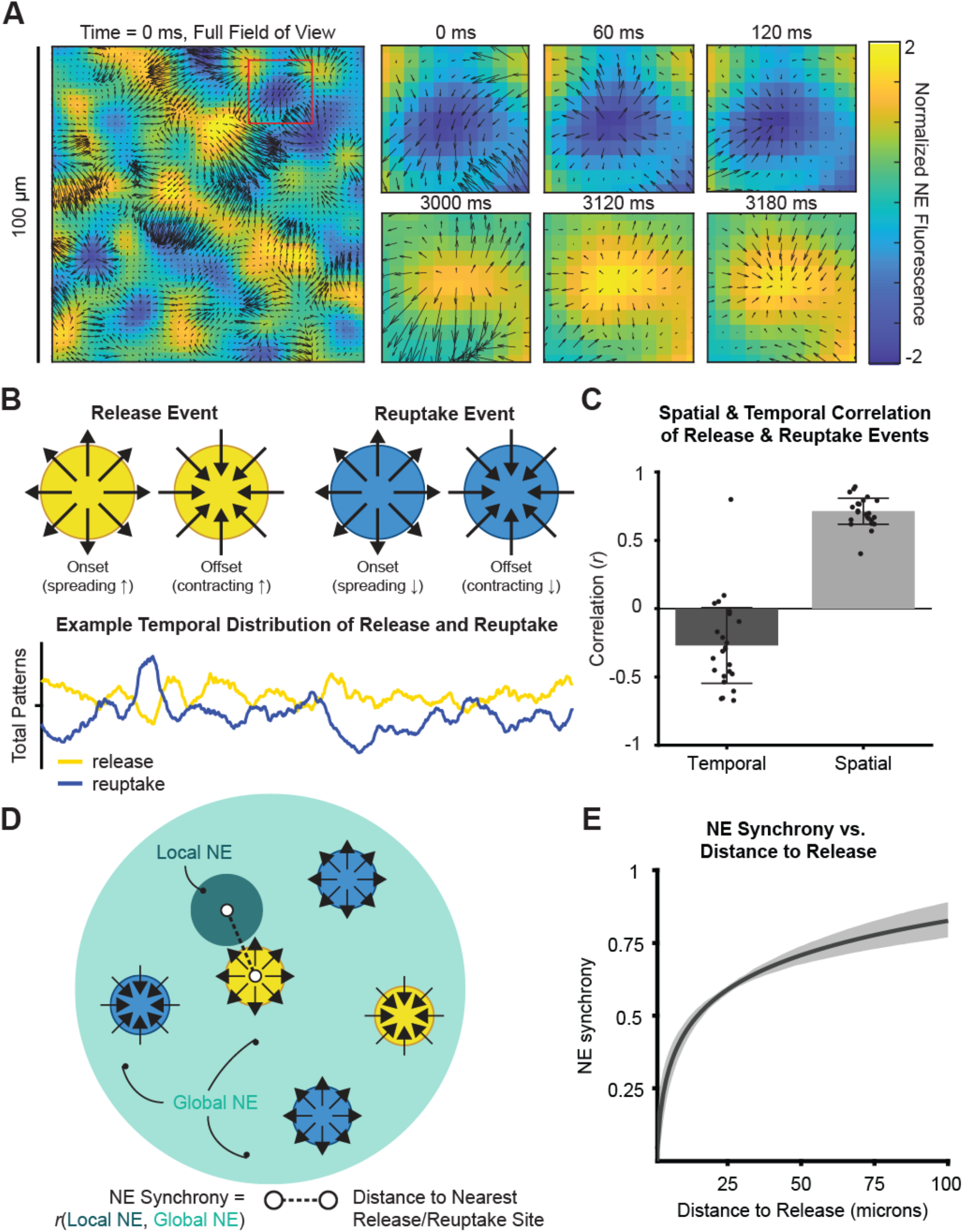
Optic flow fields reveal spatial organization of local NE. **A)** Optic flow vector fields are applied to NE fluorescence across the field of view. The spatial gradient of NE spread is used to define critical points at different time points. We observe that the same location can go from being a region of reuptake (expanding and contract region of low NE concentration) to a region of release (expanding and contracting region of high NE concentration). B) Schematic depiction of a putative release and reuptake site (top) and representative example output (bottom) from optic flow analysis of a two-photon imaging session demonstrating the number of release and reuptake sites at each point in time. C) Release and reuptake sites are spatially correlated (1-sample t-test, p = 3.2222e-20) and temporally anti-correlated (1-sample t-test, p = 5.2773e-04) across 5 minute imaging sessions (n = 24) D) Schematic depiction of how NE synchrony (sliding correlation of local and global NE in 30s windows) is related to distance to release. Distance to release is computed separately for each local NE field and is the distance (in microns) to the closest release event to that local field at each time point. E) Logistic regression results predicting NE synchrony with distance to release demonstrating asynchrony close to local NE release sites. Models were fitted for the NE synchrony time series of n = 326 local NE fields. Coefficients were significant to the alpha = 0.05 significance level in 70% of models.

## Discussion

We found that local NE^37^ is highly coupled to global NE, except during nearby release events. Local NE around a neuron improves prediction of that cell’s neural activity relative to region-level NE, particularly during periods of greater spatial variability in NE release. These relationships are disrupted by pharmacological blockade of NET, further affirming a role for spatially precise NE release. Methodologically, we introduce optic flow analyses^34^ to spatial neurotransmitter release studies which may permit the extraction of putative release and reuptake sites from two-photon neuromodulator GPCR-based sensor data. These spatial patterns are also disrupted by NET blockade. Thus, we conclude that local NE dynamics occur on a cellular scale and correlate with the activity of neural ensembles in the prefrontal cortex.

This observation raises important questions about the mechanism and function of local NE release. While the optic flow analysis we used identifies locations where NE concentration precedes NE increases elsewhere, we cannot precisely map local increases in NE to the axonal varicosities^9^ where release occurs. One prior study using a fluorescent false neurotransmitter probe for NE^14^ has identified varicosities where NE release from vesicles occurs in vivo (as well as silent varicosities where there is no release). Further in vivo imaging studies linking the structure of LC_NE_ axons near local NE release dynamics will more precisely define this relationship. Spatiotemporally precise optic control of NE release at single varicosities may also be used in concert with GPCR-based fluorescent sensors^19^ to define physiological release mechanisms. While we observed consistent and spatially localized relationships between NE and neural Ca^2+^ signals, further studies may identify cell-type or region-specific heterogeneity in this relationship.

We demonstrated that local NE around prefrontal cortical neurons influences their activity dynamics during unstructured spontaneous activity. Prior studies have demonstrated distinct changes in neural activity during task performance, such as reduced variability. Local neuromodulation has been hypothesized to subserve cognitive functions such as cognitive flexibility. Our study only addressed the release of NE during spontaneous behavior, so further studies may identify whether local NE influences on neural ensembles participates in higher order cognitive functions. Recent studies of LC-NE neurons have demonstrated that individual cells tend to fire synchronously to aversive outcomes but asynchronously to appetitive outcomes. Further studies are needed to identify the functional impact of local NE release in cognition and emotional processing.

More broadly, these results open new questions about the spatial scale of neuromodulatory processes. Spatially localized GPCR activation signals have been widely observed, but tools for analyzing those patterns require further development. The methodology we employed was to combine optic flow analysis of the GPCR fluorescent signal with GLM modeling to distinguish local release patterns and their relationships to neural activity and full-field GPCR signals. The development of spectrally orthogonal GPCR-based sensors would also permit simultaneous mapping of these GPCR release patterns. Neurons integrate influences from distinct neuromodulators in their endogenous patterns of release. Advances in GPCR-based sensors thus require advances in computational methods for defining these patterns from optic imaging data.

A neuron appears to be influenced by regional fluctuations in NE release that are distinct from other neighboring neurons. Local neighborhoods of neuromodulator concentration occur spontaneously and may affect neural activity. Our study shows that optic flow analysis applied to GPCR-based sensors can identify spatiotemporal patterns which may be related to molecular release events. Future studies can utilize this computational toolset to identify neurotransmitter release and reuptake patterns and relate these patterns to structure and function. The advent of GPCR-based fluorescence sensors has revolutionized our understanding of how neuromodulators and neuropeptides influence circuit function. Answering the question of how neurotransmitter dynamics relate to neuronal patterns of structure and function will require further studies of the precise spatiotemporal relationships between molecular signaling events in relation to neural dynamics.

## Acknowledgements, funding, and financial disclosures

We thank Jocelyne Rondeau for her help with editing this paper and making figures. We are also grateful to the funding supplied by grant R21MH118596 awarded to Dr. Kwan; the National Institutes of Health grant K08MH122733 awarded to Dr. Kaye; the Glenn H. Greenberg Fund for Stress and Resilience awarded to Dr. Kaye; the Brain and Behavior Research Foundation NARSAD Young Investigator Grant awarded to Dr. Kaye; funding from the Connecticut Mental Health Center and the National Center for PTSD granted to Dr. Kaye; and the Yale University Summer Fellowship awarded to Samira Glaeser-Khan. No conflicts of interest are relevant to the current study. Dr. Kaye receives or has received research funding from Transcend Therapeutics and Freedom Biosciences, no conflicts of interest are relevant to the current study.

